# Autophagy prevents ER stress-induced Tight Junction barrier disruption via claudin-2 homeostasis

**DOI:** 10.64898/2026.07.15.738672

**Authors:** Priya Arumugam, Kushal Saha, Ashwinkumar Subramenium Ganapathy, Alexandra Wang, Leonard Harris, Gregory Yochum, Prashant Nighot

## Abstract

Defective intestinal epithelial tight junction (TJ) barrier function and endoplasmic reticulum (ER) stress are central pathological features of inflammatory bowel disease (IBD), yet the molecular mechanisms ER stress to TJ disruption remains poorly understood. Here, we investigated the role of autophagy in regulating intestinal TJ homeostasis during ER stress. ER stress was elevated in inflamed Crohn’s disease tissue and chronic dextran sulfate sodium (DSS) colitis. In human intestinal epithelial Caco-2 monolayers, murine colon, and human colonic explants, induction of ER stress with tunicamycin, thapsigargin, or brefeldin A disrupted TJ barrier integrity, as demonstrated by reduced transepithelial electrical resistance and increased paracellular permeability. ER stress selectively increased the pore-forming TJ protein claudin-2 and altered occludin localization without significantly affecting other claudins. Pharmacologic activation of autophagy with rapamycin attenuated ER stress, restored TJ barrier function, reduced claudin-2 accumulation, and preserved occludin localization. Conversely, CRISPR-Cas9-mediated deletion of autophagy gene ATG7 exacerbated ER stress, apoptosis, and TJ barrier dysfunction in vitro, while intestinal epithelial-specific Atg7 knockout mice exhibited enhanced ER stress-induced intestinal permeability in-vivo. Mechanistically, prolonged ER stress impaired autophagic flux through IRE1α kinase signaling, resulting in accumulation of p62 and claudin-2. Inhibition of IRE1α kinase activity restored autophagy, reduced claudin-2 levels, and preserved TJ barrier function. We further identified adaptor-associated kinase 1 (AAK1) as a downstream mediator of IRE1α signaling during ER stress, with increased AP2M1 phosphorylation and altered claudin-2 trafficking. Claudin-2 overexpression alone induced ER stress and lysosomal damage, suggesting a feed-forward mechanism amplifying epithelial injury. Finally, enteric rapamycin administration reduced ER stress and restored autophagy in murine DSS colitis. Collectively, these findings identify an IRE1α–AAK1–autophagy axis as a critical regulator of intestinal TJ barrier integrity during ER stress.

## Introduction

The apically located inter-cellular tight junctions (TJ) polarize the intestinal epithelial cell into apical and basolateral regions (fence function) and forms a selective barrier against paracellular permeation of noxious luminal antigens (gate function)^1, 2^. Defective intestinal TJ barrier is an important pathogenic factor for IBD, particularly Crohn’s disease (CD) and ulcerative colitis (UC) which together cost the U.S. more than $8 billion annually^3–5^. The defects in intestinal TJ barrier allow increased antigenic penetration and amplify inflammatory response in intestinal tissue^4, 6^. The progress in therapeutic modulation of TJ barrier function, however, has been minimal.

TJs consist of membrane-spanning proteins (e.g., occludin and claudins) linked by cytoplasmic plaque proteins, zona occludin-1 and −2, to the cytoskeleton. Functionally, the TJs are known to have a small size, cation selective “pore” pathway and a large size, non-charge selective “leak” pathway. The pore pathway is defined by the channel-forming claudin-2 while occludin is involved in the regulation of the leak pathway^7, 8^. Recently, we have defined the role of autophagy in promoting the intestinal epithelial TJ barrier by regulation of intracellular trafficking of TJ proteins^9–12^.

Autophagy (macroautophagy) is an intracellular degradation system that delivers diverse cellular cargo sequestered inside double-membrane vesicles (autophagosomes) to the lysosome. In a well-regulated, multi-step process, several autophagy-related (ATG) proteins take part in cargo sequestration and autophagosome formation. Genome-wide association studies have identified mutations in ATG genes as risk factors for IBD^13–15^. Recent studies have shown the role of autophagy in dendritic-epithelial cell interactions, NOD2-directed bacterial sensing, and immune-mediated pathogen clearance^16–19^. In clinical IBD studies, persistent increase in intestinal permeability predicts poor clinical outcome, and normalization of intestinal permeability correlates with long-term clinical remission^20, 21^. Similarly, in animal models of IBD, enhancement of intestinal TJ barrier ameliorates intestinal inflammation^22, 23^. In line with these observations, we have shown that autophagy differentially regulates TJ proteins to enhance TJ barrier; it degrades channel-forming claudin-2 and increases levels and TJ localization of barrier-forming occludin^10–12, 24^.

Protein folding occurs in the endoplasmic reticulum (ER) prior to transport to their intracellular or extracellular destinations. Under cellular stress conditions, unfolded or misfolded proteins accumulate in the ER lumen, triggering the unfolded protein response (UPR) to restore homeostasis. The UPR mainly involves three distinct components: transcriptional induction of ER-resident chaperone genes to facilitate protein folding, translational attenuation to decrease the demands made on the organelle, and ER-associated degradation (ERAD) of the unfolded proteins^25–28^. Autophagy is thought to be an alternate way of degradation of misfolded proteins^29^. Three major pathways activating transcription factor-6 (ATF6), inositol-requiring enzyme 1 (IRE1), and double-stranded RNA-dependent protein kinase (PKR)-like eukaryotic initiation factor 2α (eIF2α) kinase (PERK) constitute UPR. IRE1 is the most evolutionally conserved and possesses both Serine/Threonine-Protein Kinase and endoribonuclease activities; it activates X-box binding protein mRNA (XBP) 1 by splicing which in turn regulates several cell survival genes^26^. Among two isoforms, IRE1α promotes while IRE1β protects from intestinal inflammation.^30^ Thus, ER stress is a common denominator of various pathophysiologic processes and diseases including IBD^31, 32^.

The intestinal epithelial cells in IBD and other intestinal diseases display excessive ER stress^33–36^. Moreover, multiple mouse models have shown a causative role for ER stress in intestinal inflammation^37–40^. Also, inhibition of ER stress can ameliorate experimental colitis^38, 39, 41, 42^. However, there is a crucial gap in our knowledge regarding the ER stress mechanisms that lead to disruption of the TJ barrier. Our study shows that autophagy plays a critical role in the preservation of intestinal TJ barrier during the ER stress via claudin-2 homeostasis.

## Materials and Methods

### Chemicals and antibodies

Rapamycin (Life Technologies, PH21235), Tunicamycin (Cell signaling, 12819S), Brefeldin A (Biolegend, 420601), Thapsigargin (Fisher Scientific, NC1412406), GSK2850163 (Sigma Aldrich, SML1684) were purchased from the indicated companies. [^3^H] Inulin (specific activity 450 mCi/g) was purchased from American Radiolabeled Chemicals, Inc. The primary antibodies used included anti-CLDN1, anti-CLDN3, anti-CLDN4, anti-phospho IRE1α (Invitrogen, 37-4900, 34-1700, PA5-34437, PA1-16927, respectively), anti-CLDN2 (Abcam, ab53032), anti-LC3, anti-ATG7 (Sigma Aldrich, L7543, A2856, respectively), anti-phospho-AP2M1, anti-CHOP, anti-IRE1α, anti-Cathepsin B (Cell signaling Technologies, 7399S, 2895, 3294S, 31718), anti-BIP, anti-SQSTM1/p62, anti-ACTB/β-actin (Protein tech, 66574-1, 18420-1-AP, HRP-60008, respectively). The secondary antibodies included anti-rabbit HRP (Invitrogen, 31460) and anti-mouse HRP (Invitrogen, 31430). TaqMan-tagged primers for assaying the expression of all the target genes by qRT-PCR were purchased from Life Technologies.

### Cell Culture

Human intestinal epithelial Caco-2 cells (ATCC, HTB-37) were maintained in DMEM (Dulbecco’s Modified Eagle’s Medium) – High Glucose (Gibco, 11965118) supplemented with 10% heat inactivated fetal bovine serum (R&D systems, S11150H) and Penicillin-Streptomycin antibiotics (Gibco, 15140122) at 37°C in a 5% CO_2_ incubator. Caco-2 cell monolayers were grown on 12 mm transwell with 0.4 μm pore polyester membrane inserts (Corning, 3460). The trans-epithelial electrical resistance (TER) of the filter-grown cells was measured by an epithelial voltohmeter and STX2 electrode set (World Precision Instruments, Sarasota, FL, USA), and monolayers with a TER of 450–500 Ω/cm^2^ were used for experiments. Cells in standard growth medium were treated with Tunicamycin (Tn, 20 μM), rapamycin (1 μM), Brefeldin A (0.5X), thapsigargin (0.8 μM), GSk2850163 (20 nM) for 24 h or indicated otherwise.

### Determination of Caco-2 Paracellular Flux

Caco-2 paracellular permeability was determined using the paracellular marker Inulin (^3^H, M_r_ = 60). The apical-to-basal flux rates of the paracellular marker was determined by adding them to the apical solution and radioactivity was measured in the basal solution at 30 min and 60 min using a scintillation counter, as described by us previously ^43^.

### Western Blot Analysis for Assessment of Protein Expression

Caco-2 monolayers were rinsed twice with ice-cold PBS (Corning, 21-040-CV) and lysed using RIPA buffer (Sigma, R0278) containing complete mini, protease inhibitor (Sigma, 11836170001). The mouse colonic mucosa or human colonic mucosa were washed with ice-cold PBS and snap-frozen in liquid nitrogen soon after harvesting or treatment. The frozen tissues were resuspended in RIPA buffer containing protease inhibitors, minced and sonicated to prepare tissue lysates. The cell/tissue lysates were centrifuged at 10,000 rpm, for 10 min in order to remove cell debris and the clear supernatant was used for protein quantification. Protein quantification of the extracted aliquots was performed (BCA protein assay kit; Pierce, 23225), and Laemmli gel loading buffer (Invitrogen, NP007) was added to the lysate and boiled at 70°C for 10 min. An equal concentration of protein was loaded in SDS-PAGE gel, separated, and transferred to a nitrocellulose membrane. The membrane was incubated for 1 h in blocking solution (5% non-fat dry milk (Bio-Rad, 1706404) in TBS (Bio-Rad, 1706435)-0.1% Tween 20 (Bio-Rad, 1706531) buffer, followed by incubation with the appropriate primary antibody in blocking solution. After incubation with primary antibody, the membrane was washed in TBS-0.1% Tween 20 buffer, incubated in the appropriate secondary antibody and developed using SuperSignal West Pico PLUS kit (Thermo Scientific, 34580). The densitometry analysis was performed using ImageJ software^44^.

### Confocal immunofluorescence

Confocal Immunofluorescence for CLDN2, and Ocln and cathepsin B on Caco-2 cell monolayers was performed by standard methods. Caco-2 monolayers were washed twice with cold PBS, fixed with 2% paraformaldehyde for 20 min. The cell monolayers were permeabilized with 0.1% Triton X-100 (Sigma, X100) in PBS at room temperature for 5 min. The cell monolayers were then blocked in normal serum (Invitrogen, 50197Z) and labeled with primary antibodies in blocking solution overnight at 4°C. After PBS washes, the sections were incubated in Alexa Fluor-488, Cy-3, or Alexa Fluor-647-conjugated secondary antibodies (Invitrogen, A11078, A10521, and A21244). ProLong Gold antifade reagent (Invitrogen, P36931) containing DAPI as a nuclear stain was used to mount the sections on glass slides. The slides were examined using a confocal fluorescence microscope Leica SP8 (Penn State College of Medicine, Advanced Light Microscopy Core, RRID:SCR_022526). Images were processed with LAS X software (Leica Microsystems,).

### Lentiviral mediated CRISPR-Cas9 knockout of *ATG7*

The plasmid Cas9 nuclease CP-LVC9NU (Genecopoeia) was used individually with a single guide RNA (sgRNA) targeting the region AAATAATGGCGGCAGCTACG of *ATG7* and scrambled sgRNA for control in pCRISPR-LVSG03 (Genecopoeia) to generate respective lentiviral particles packaged in Lenti-X 293T cells using PMD2.G and pPAX2 (Addgene, 12259, 12260; deposited by Didier Trono). The lentiviral particles obtained were used to transduce Caco-2 cells in presence of polybrene (EMD Millipore, TR-1003-G) and were selected in their respective antibiotic selection media to generate stable knockout Caco-2 cells. The gene knockout was further confirmed using western blot analysis. CLDN2 ORF in pCMV6-AC-GFP (Origene, RG204199) and corresponding control plasmid were used to transfect Caco-2 cells using Lipofectamine 2000 (Invitrogen, 11668027) as per manufacturer’s instructions and the transfected cells were selected using respective antibiotic selection media to generate stable CLDN2-overexpressing Caco-2 cells.

### Experimental animals

Experimental methodologies used in the study were approved by the Institutional Animal Care and Use Committee of Pennsylvania State University College of Medicine. Adult mice were engineered with floxed alleles of *Atg7* and a transgene expressing the TAM-regulated Cre recombinase fusion protein under the control of the intestinal epithelial cell specific *Villin* promoter (Vil-CreERT2)^45^. ATG7 deficiency was created by providing tamoxifen to 10-week-old mice leading to Cre activation only in mice with *Atg7* floxed and Vil1-Cre/ERT2alleles, producing intestinal epithelial specific loss of ATG7 protein (Atg7*^IECΔ/Δ^* mice). Tamoxifen-treated mice with only floxed alleles of *Atg7* (*Atg7*^fl/fl^) were used as controls. Tamoxifen (Sigma, T5648; 20 mg/ml suspended in 98% sunflower seed oil [Spectrum Chemical, S1929] and 2% ethanol mixture) was injected i. p. (200 µl per 25 g of mice body weight) into 8 to 10 week old Vil1-cre/ERT2^+/+^ *Atg7*^fl/fl^ mice once per day for 5 days and the mice were used for experiment after 2 weeks. Control animals *Atg7*^fl/fl^ mice also received the same amount of tamoxifen. In the chronic dextran sulfate sodium (DSS) model, mice were administered 1.5% DSS (MP Biomedicals, 160110) in drinking water for 5 days, followed by 5 days of regular water, repeated for three cycles. Microencapsulated rapamycin (eRapa diet) (or diet with capsule material only) was obtained from Rapamycin Holdings, San Antonio, Tx, formulated into 5LG6 diet by Purina, and supplied by LabDiet.

### Human Tissue samples and treatment

The surgically resected human colon samples were obtained freshly from the Department of Surgery, Division of Colon and Rectal Surgery as per the protocols approved by Institutional Review Board (STUDY00010256). Human colonic tissue were washed, the muscular layer was stripped, and the isolated mucosal epithelial tissue were incubated in supplemented DMEM, overnight, on gelatin sponge (Ethicon, 1975), in the presence and absence of Tunicamycin (10 µg/ml).

### Measurement of paracellular permeability and trans-epithelial electrical resistance (TER) of murine and human colon

The trans-epithelial resistance of the mice and human colonic tissue were measured by mounting the colonic tissue on 0.03 cm^2^-aperture Ussing chambers (Physiologic Instruments, CA, USA). Trans-epithelial electrical resistance (TER, Ω·cm^2^) was calculated from the spontaneous potential difference and short-circuit current. The paracellular permeability was assessed by mucosal-to-serosal flux of [^14^C]-urea, as described by us previously ^46^.

### Cell transfections

hIRE1α K599A was a gift from Fumihiko Urano (Addgene plasmid #20745; RRID). The human IRE1α kinase mutant K599A expressing plasmid was transiently transfected into WT and IRE1-KO Caco-2 cells using Lipofectamine 3000 (Thermo Fisher Scientific, L3000015). Briefly, plasmid DNA was diluted in Opti-MEM medium. Lipofectamine 3000 reagent was diluted separately in Opti-MEM and mixed with the diluted DNA solution for 15 min at room temperature to allow complex formation. The DNA-lipid complexes were then added to the cells and incubated under standard culture conditions. Cells transfected with the corresponding empty vector served as controls. Following transfection, cells were incubated for 24 h. Expression of IRE1α K599A was confirmed by immunoblotting analysis. For ectopic expression of Cldn2, Caco-2 cells were transfected with a Cldn2 expression clone (Ex-V1294, vector pEZ-Mo2, GeneCopoeia), according to the instructions of the supplier.

### NanoBret target engagement assay

Intracellular AAK1 target engagement was assessed using the NanoBret® Target engagement intracellular kinase assay according to the manufacturer’s instructions. Scrambled control and IRE1Δ Caco-2 cells were transiently transfected with a NanoLuc®-AAK1 fusion construct and seeded into 96-well tissue culture treated white plates. Following overnight incubation, cells were treated with Tn (20 μM) or vehicle control for 24 h. NanoBret® kinase tracer was added to the cells and incubated to allow intracellular binding to the AAK1 kinase domain. Subsequently, NanoBret® NanoGlow® substrate were added according to manufacturer’s protocol. Luminescence was measured at donor (460nm) and acceptor (610 nm) wavelengths using a spectrophotometer. BRET ratios were calculated.

### Statistical analysis

Data are reported as means ± SE. Whenever needed, data were analyzed by using an ANOVA for repeated measures (SigmaStat, Systat Software, San Jose, CA). Tukey’s test was used for post-hoc analysis between treatments following ANOVA (*P* < 0.05).

## Results

### ER stress is a fundamental element of IBD

Elevated ER stress has been shown to be a pathogenic factor in IBD ^34–36^ as well as several IBD models^25, 35, 36, 38, 40^. Corroborating previous studies, we found increased ER stress in CD inflamed tissue compared to adjoining non-inflamed tissue in paired ileal samples from CD patients (**Fig. 1**) and chronic dextran sodium sulfate (DSS) model of IBD (**Fig. 9D**).

**Fig. 1.**
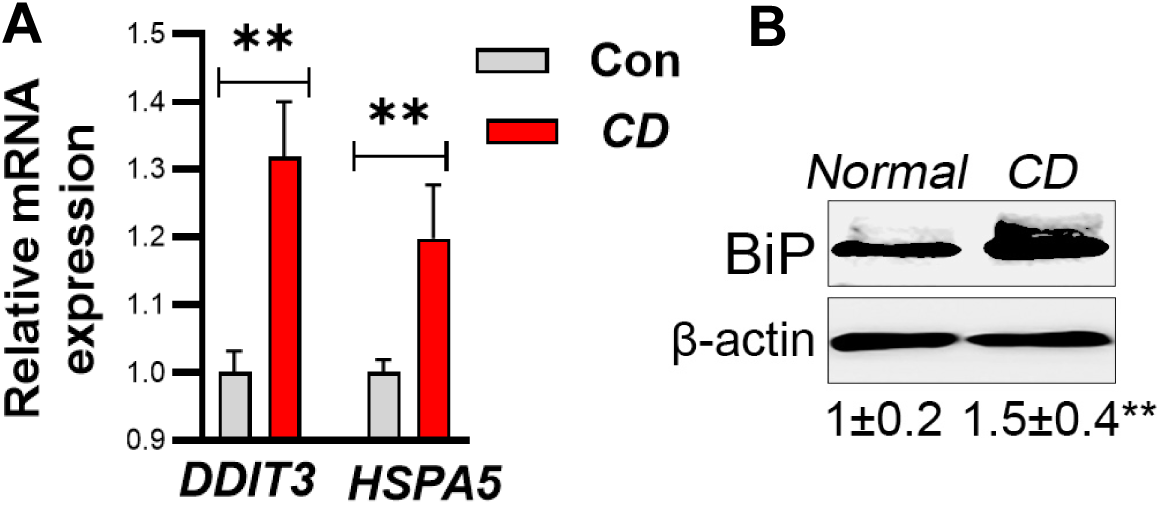
ER stress in Crohn’s disease. A) Compared to normal adjacent tissue, moderately inflammed Crohn’s disease (CD) ileal tissue showed increased expression of UPR genes DDIT3 (CHOP) and HSPA5 (BiP). (B) CD tissue also showed increased protein levels of BiP. The numbers below the bands indicate densitometry. **, p<0.01.

### ER stress induces intestinal TJ barrier disruption

To study the effect of ER stress on intestinal epithelial TJ barrier function, we treated filter-grown human intestinal Caco-2 cell monolayers, which form a robust TJ barrier, with tunicamycin, a well-known ER stress inducing agent that blocks protein N-linked glycosylation. Tunicamycin-induced ER stress was evident by increased expression of UPR genes including C/EBP homologous protein (CHOP), glucose-regulated protein (GRP78 or BiP), and activating transcription factor-4 (ATF4) (**Fig. 2A**) and proteins (**Fig. 2B**). Tunicamycin caused a reversible drop in the transepithelial resistance (TER) (**Fig. 2C**) and caused an increased flux of paracellular probes inulin (**Fig. 2D**), urea and 4k dextran (not shown). Thus, tunicamycin disrupted TJ barrier in Caco-2 monolayers. Other ER stress inducers thapsigargin and brefeldin caused a similar reduction in the Caco-2 TJ barrier (**Fig. 2E**). Tunicamycin treatment disrupted TJ barrier in other cell lines including T84 and non-transformed model epithelia MDCKII. Tunicamycin also induced ER stress and disrupted colonic TJ barrier in wild type (WT) mice, as reflected by the decreased TER and increased inulin flux in Ussing chamber studies (**Fig. 3A, B,** and **C**). Moreover, in healthy human colonic explant culture, tunicamycin induced ER stress, reduced the TER, and increased the paracellular permeability (**Fig. 3D, E,** and **F**).

**Fig. 2.**
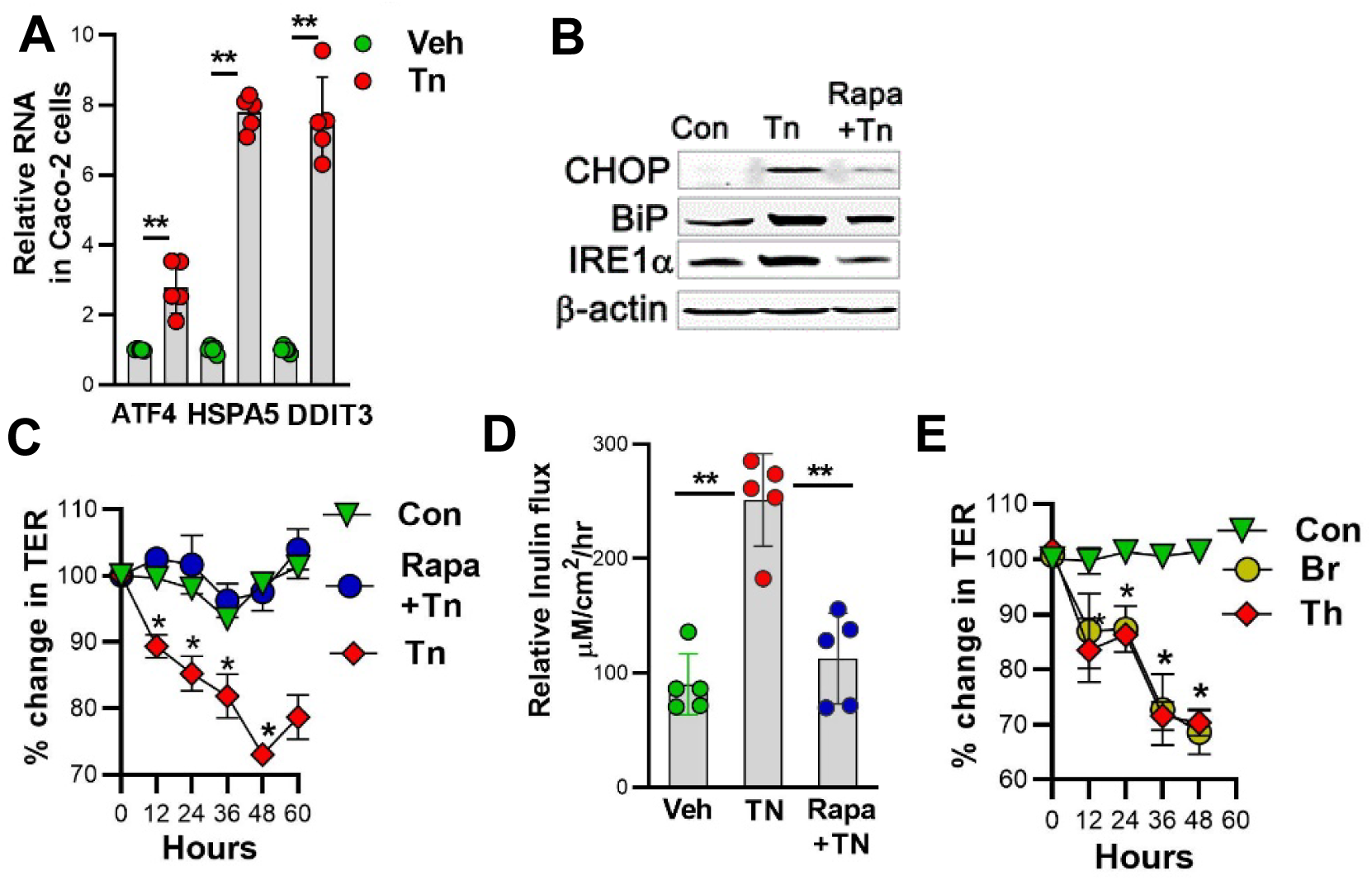
Tunicamycin induces ER stress and TJ barrier dysfunction in Caco-2 cells. Tunicamycin (Tn), 10mg/ml, 24 hrs, increased expression of UPR genes Activating transcription factor-4 (ATF4), C/EBP homologous protein (CHOP), and glucose-regulated protein (GRP78 or BiP). (B) Tn also increased ER stress proteins CHOP, BiP, and IRE1a. Tn reduced transepithelial resistance (TER) (Tn media changed at 48 hrs) (C), and increased paracellular inulin (D). Thapsigargin (Tg) and brefeldin (Br) also caused reduction in Caco-2 TER (E). Rapamycin (500nm, Rapa) prevented Tn-induced increase in UPR protein expression (B) and disruption of TJ barrier (C and D). **, p<0.01*, p<0.05 vs vehicle.

**Fig. 3.**
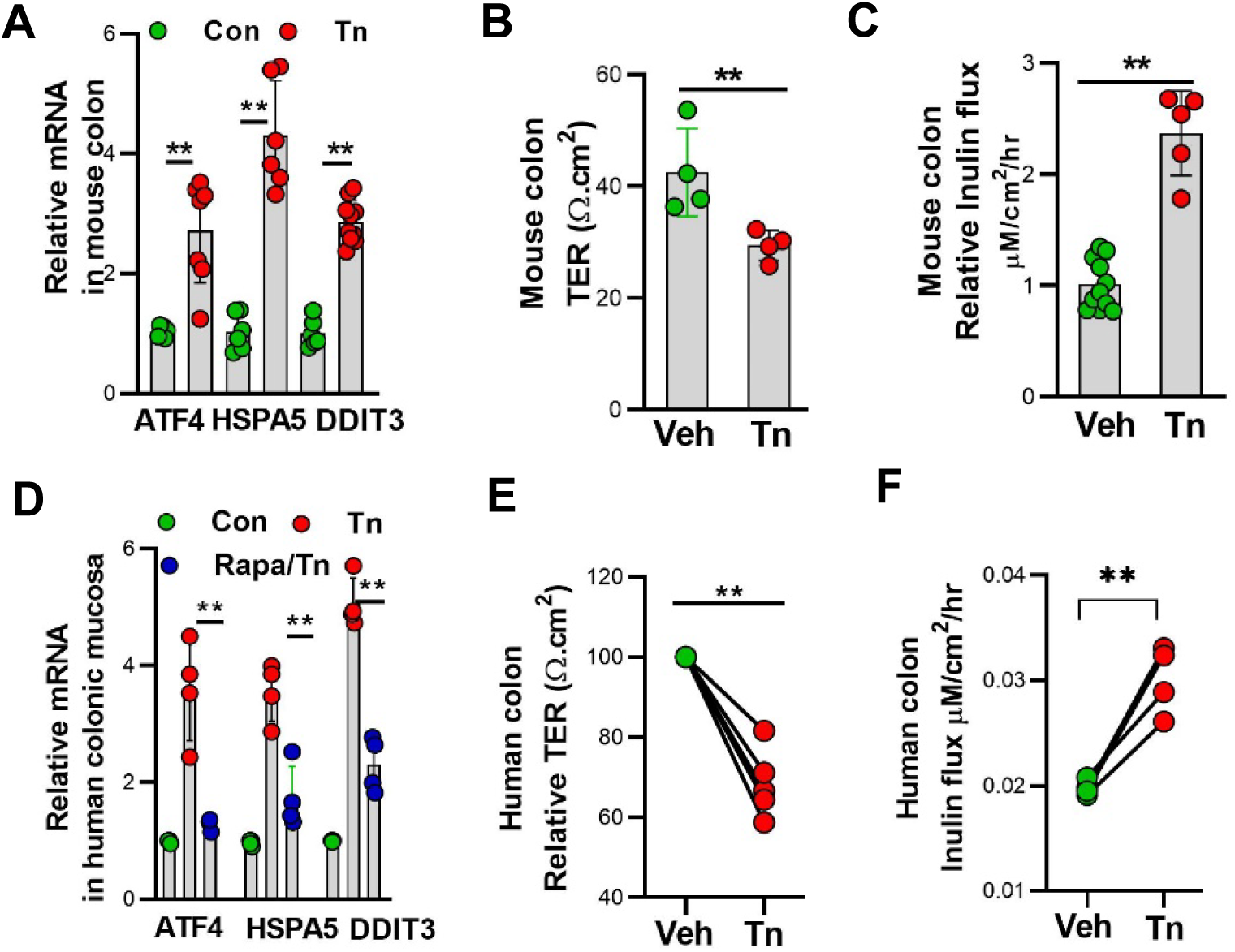
ER stress induces TJ barrier dysfunction in mouse and human colon. (A) Tunicamycin (Tn, 1mg/kg, i.p.) induced ER stress and increased UPR gene expression (ATF4, HSPA5, and DDIT3) in mouse colonic mucosa. Tn reduced TER (B), and increased paracellular inulin flux (C) in mouse colons. Mouse colons were mounted in Ussing chambers after 24 hours of Tn administration. Tn (10μg/ml, 18 hours) induced UPR genes expression (D), reduced TER (E), and increased paracellular inulin flux (F) in human colonic explants. Rapamycin (Rapa) significantly prevented Tn-induced UPR gene expression in mouse colon (A) and human colonic explants (D). Rapamycin was injected i.p. (250 μg/kg) in mice. Paired human healthy colonic mucosa was cultured using gelatin sponge for 18 hours with Tn or vehicle and mounted in Ussing chambers. **, p<0.01 vs vehicle control.

### ER stress increases channel-forming TJ protein claudin-2 levels

In view of the TJ “pore” and “leak” pathways being regulated by specific proteins^7, 8^, we examined the TJ composition after tunicamycin treatment. Among the select claudins known to regulate paracellular TJ barrier^47^, the protein level or localization of claudin-1, −3, −4 and occludin showed no significant change after tunicamycin treatment. However, tunicamycin markedly increased the levels of channel-forming claudin-2 and led to its cytoplasmic accumulation in Caco-2 cells (**Fig. 4A, B, and C**). We also examined the TJ composition after tunicamycin treatment of human colonic explants from normal tissue. Among the select claudins regulating the TJ barrier^47^, tunicamycin markedly increased the levels of channel-forming claudin-2 (**Fig. 4D)**. Thus, ER stress specifically causes an increase in claudin-2 levels. The mRNA levels of claudin-2 and occludin were not changed significantly following the tunicamycin treatment.

**Fig. 4.**
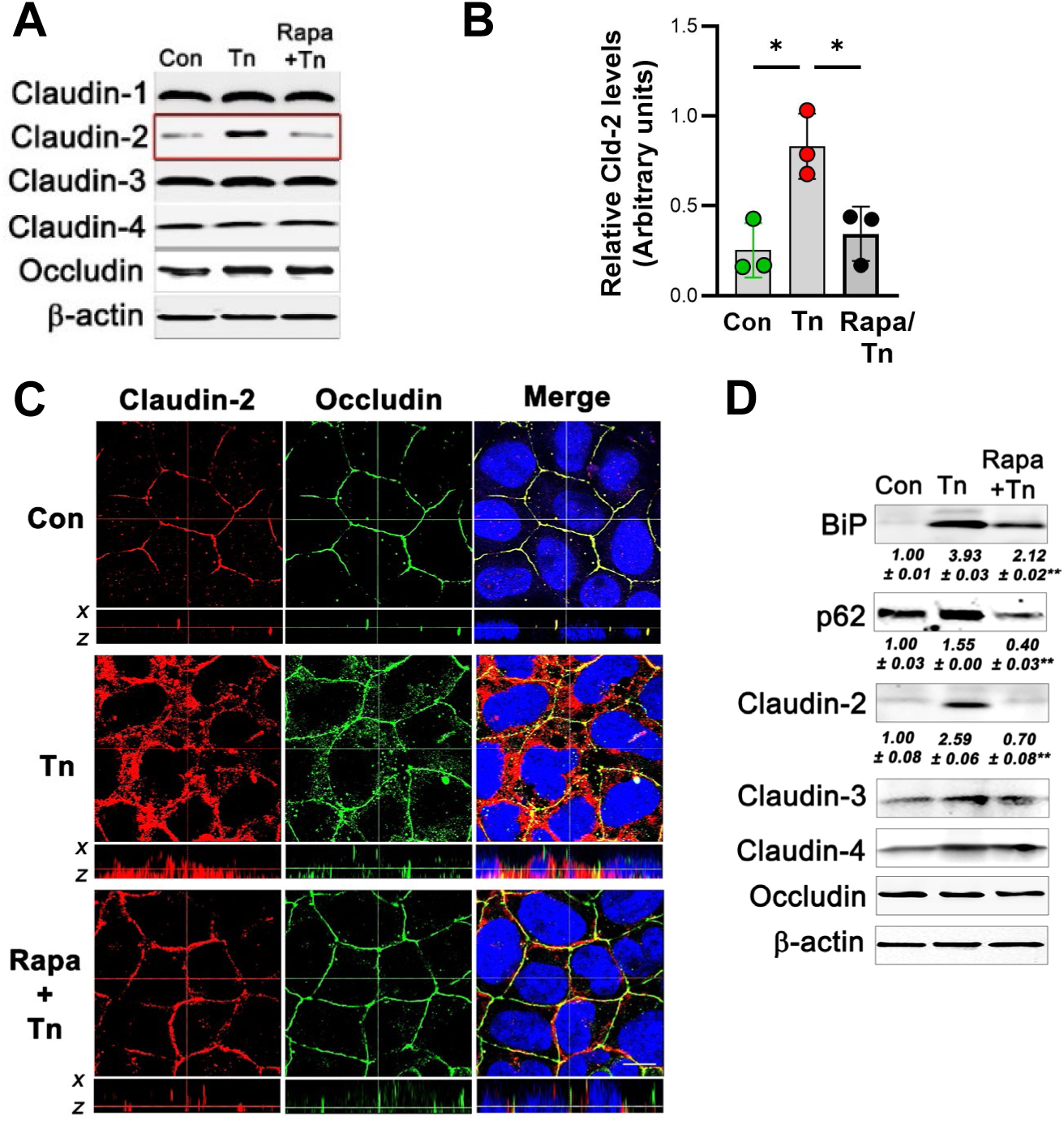
Tunicamycin (Tn) increases claudin-2 levels. (A) Tn increased claudin-2 levels in Caco-2 cells. Tn: 10μg/ml, 24 hrs. n = 3 blots. (B) Densitometry for claudin-2 bands in panel (A). (C) Claudin-2 (red) and occludin (green) were redistributed to cytoplasm by Tn treatment. Rapa largely prevented Tn-induced increase in cytoplasmic claudin-2 and occludin redistribution. The X-Z planes showed that Rapa restored levels and localization of claudin-2 to the apical TJ puncta after ER stress. The dotted lines represent optic level in each panel. White bar = 5μm. (D) Tn induced ER stress (increased BiP levels), inhibited autophagy (reduced p62 levels) and increased claudin-2 levels in human colonic explants. Tn: 10μg/ml, 12 hrs. Rapamycin (Rapa) reversed Tn effect. The numbers below the BiP, p62 and claudin-2 blots indicates densitometry. **, p< 0.001.

### Autophagy is protective against ER stress-associated TJ barrier disruption

As autophagy is cytoprotective^48^, we examined the role of autophagy in ameliorating ER stress-induced TJ barrier disruption. Rapamycin, an inhibitor of mTOR and activator of autophagy^49^, dampened the tunicamycin-induced ER stress (**Fig. 2A and B**) and potently prevented tunicamycin-induced drop in TER (**Fig. 2 C**), increase in paracellular flux (**Fig. 2D**), increase in claudin-2 levels (**Fig. 4A and B**), and occludin de-localization **(Fig. 4C)** in Caco-2 cells. Rapamycin also potently prevented tunicamycin-induced ER stress (**Fig. 3D and 4D**), increase in claudin-2 levels and promoted autophagy (reduction in p62 levels) in human colonic explants **(Fig. 4D)**.

### Disruption of autophagy increases ER stress and associated TJ barrier disruption

To further validate the role of autophagy in maintaining the cell integrity during the ER stress-induced TJ barrier disruption, specific autophagy related ATG genes^50, 51^ were knocked out in Caco-2 cells using CRISPR-Cas9 method. Silencing of ATG7 (*ATG7Δ*) significantly increased tunicamycin-induced ER stress (**Fig. 5A** and **B**), and apoptosis (**Fig. 5C**). Similar findings were noted in ATG5 and ATG16 deleted Caco-2 cells (not shown). Also, tunicamycin-induced increase in TJ permeability was significantly higher in *ATG7Δ* Caco-2 cells compared to scramble cell (**Fig. 5D**). Next, we examined the intestinal epithelial specific role of autophagy against ER stress-induced TJ barrier disruption. Tunicamycin-induced increase in colonic TJ permeability was significantly higher in intestinal epithelium specific (Cre-villin) *Atg7^IECΔ/Δ^* mice compared to Atg7^fl/fl^ mice (**Fig. 5E and F**).

**Fig. 5.**
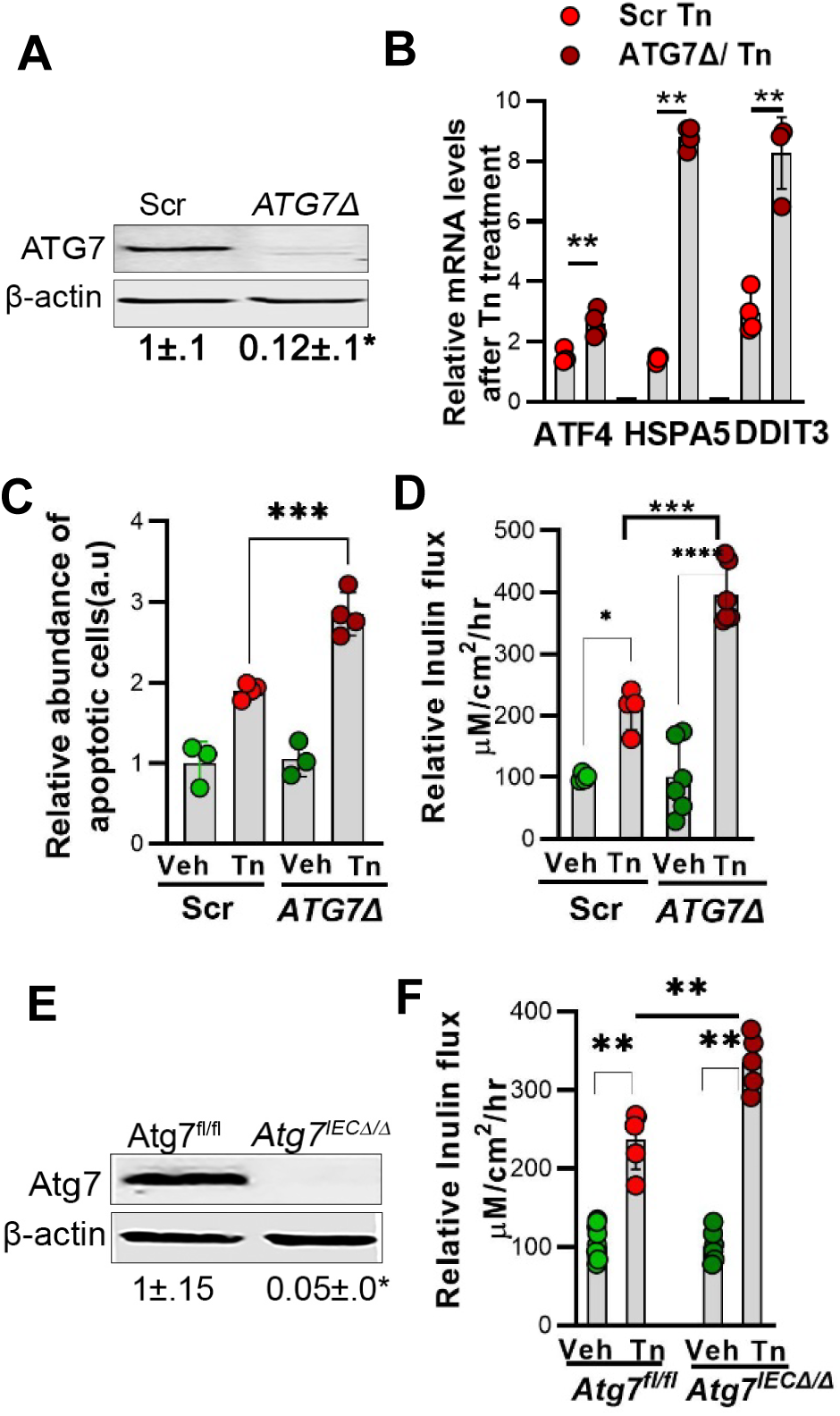
Disruption of autophagy increases ER stress and associated TJ barrier disruption. (A) CRISPR-Cas9-mediated deletion of Atg7 (ATG7Δ) in Caco-2 cells. *p< 0.03. Tn-induced ER stress (B), apoptosis (C) and TJ permeability (D) was significantly increased in ATG7Δ cells. (E) Intestinal epithelium specific (Cre-villin) knockout of Atg7 in mice (Atg7^IECΔ/Δ^). *p< 0.03. (F) Atg7^IECΔ/Δ^ mice showed exaggerated increase in inulin flux compared to Atg7^fl/fl^ mice after Tn administration. *, **, ***, p<0.001, One-way ANOVA.

### ER stress disrupts autophagy-claudin-2 homeostasis

Even though it is widely known that the IRE1-JNK pathway promotes cytoprotective autophagy during the early stages of ER stress (2-4 hours),^52^ we noted that autophagy is stifled during persistent ER stress, in-vitro and in-vivo. Caco-2 cells treated with tunicamycin showed reduction in p62 levels at 24 hours. However, the p62 levels were increased after 48 hours of tunicamycin treatment, indicating inhibition of autophagy at a later time point (**Fig. 6A**). The p62 levels were also increased in the colon of mice administered with tunicamycin after 72 hours (**Fig. 6B**), indicating inhibition of autophagy after prolonged ER stress. Next, to understand how ER stress induces TJ barrier disruption, we inhibited the three ER stress signaling pathways: PERK, IRE1, and ATF6 during tunicamycin treatment. While PERK and ATF6 inhibition had no effect on the tunicamycin-induced loss of Caco-2 TER, IRE1α kinase inhibition prevented tunicamycin-induced drop in TER (**Fig 6C**). Moreover, IRE1α kinase also prevented tunicamycin-induced increase in claudin-2 levels (**Fig. 6D**). We also observed that IRE1α kinase inhibition promoted autophagy during the ER stress, as indicated by reduction in the ER stress-induced accumulation of p62 and increase in LC3-II/-I levels (**Fig. 6D**). Thus, the dominant role of IRE1α kinase activity in ER stress-induced autophagy inhibition and increase in claudin-2 levels is suggested by the reversal of both processes by its inhibition.

**Fig. 6.**
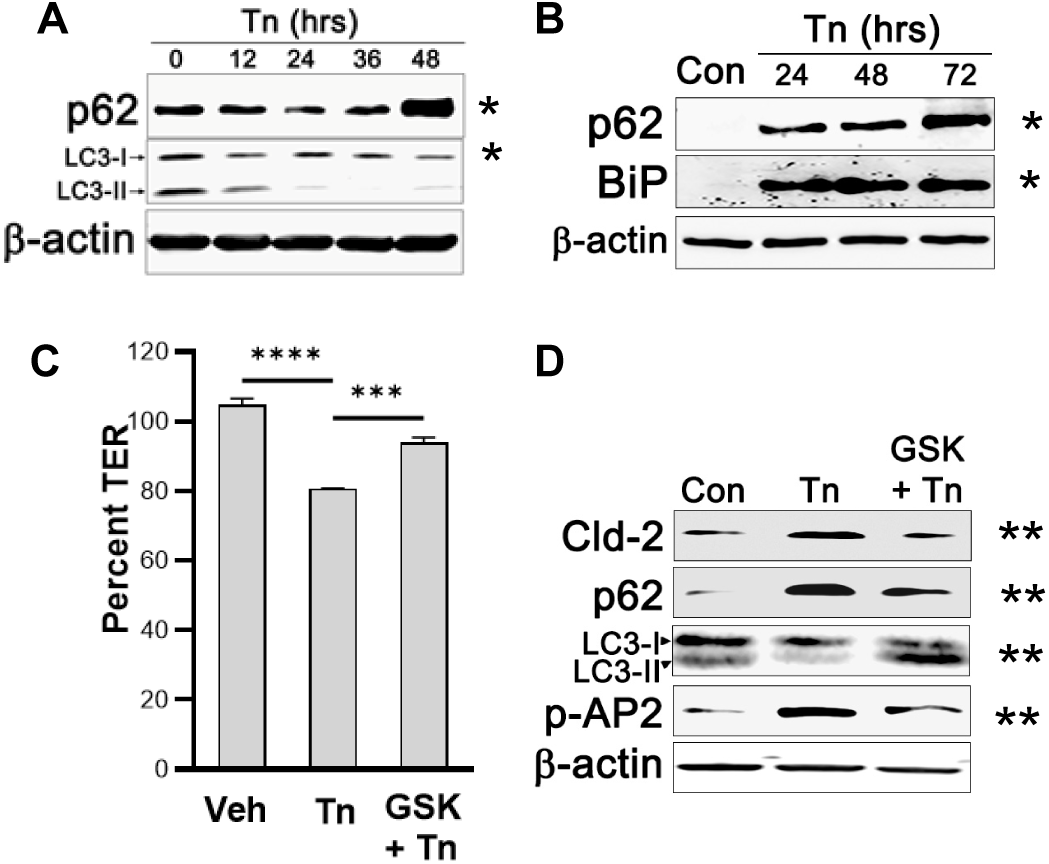
ER stress disrupts autophagy-claudin-2 homeostasis. (A) p62 levels increased and LC3-II levels decreased at 48 hours after tunicamycin (Tn) treatment, indicating inhibition of autophagy in Caco-2 cells. (B) Tn also increased p62 levels in mouse colon. (C) IRE1 kinase inhibition (GSK2850163, 20nM) prevented Tn-induced drop in TER. ***, p<0.001, ***, p<0.001. (D) IRE1 kinase inhibition (GSK2850163, 20nM) reduced Tn-induced increase in claudin-2 (Cld-2) and promoted autophagy as evident by reduction in p62 and an increase in LC3-II levels. GSK also inhibited Tn-induced activation of AP2M1 (p-AP2) in Caco-2 cells. Represents 3 blots. *, p<0.03 Con vs Tn; **, p<0.03 Tn vs GSK+Tn, densitometry by ImageJ.

Based on our previous findings of autophagy mediating the enhancement of TJ barrier via degradation of channel-forming claudin-2,^10, 24^ we hypothesize that IRE1α-mediated inhibition of autophagy plays a critical role in disruption of the TJ barrier during the ER stress. Thus, in contrast to IRE1-JNK pathway-mediated promotion of cytoprotective autophagy during the early stages of ER stress (2-4 hours),^52^ we observed that IRE1α activation during persistent ER stress abrogates autophagic process.

### IRE1 activates AAK1 to disrupt autophagy and claudin-2 homeostasis

Recently, we have discovered AAK1 as a novel regulator of the TJ barrier. We showed that the AAK1-mediated activation of adaptor protein (AP2) μ subunit AP2M1, plays a crucial role in anchoring membrane claudin-2 to clathrin and LC3, for ultimate autophagic degradation.^10^ Here, we found that AAK1 is remarkably aggregated with claudin-2 in the cytoplasm during the ER stress in Caco-2 as well as non-transformed, claudin-2 rich HIEC cells. Such an abnormal activation and accumulation of AAK1 with toxic protein aggregates has been described in ER stress-related neurodegenerative diseases.^53, 54^ We also found that the phosphorylation of only known AAK1 target AP2M1 is significantly increased during the ER stress and IRE1α kinase inhibition prevented tunicamycin-induced phosphorylation of AP2M1 (**Fig.6C**), suggesting that AAK1 may be a direct target of IRE1. Thus, we hypothesize that the IRE1α-mediated excessive activation of AAK1 during the ER stress causes cytoplasmic aggregation of claudin-2 and that degradation of cytoplasmic claudin-2 is hampered due to impaired autophagy. It is also noteworthy that over-expression of claudin-2 alone increased ER stress (**Fig. 7A and B**). As ER stress also causes lysosomal damage, we examined if claudin-2 overexpression affects lysosomes. As reflected by increased and diffuse staining of Cathepsin B, claudin-2 overexpression alone caused lysosomal damage similar to tunicamycin-induced ER stress (**Fig. 7C**) underlining the detrimental effects of increased cellular claudin-2 levels.

**Fig. 7.**
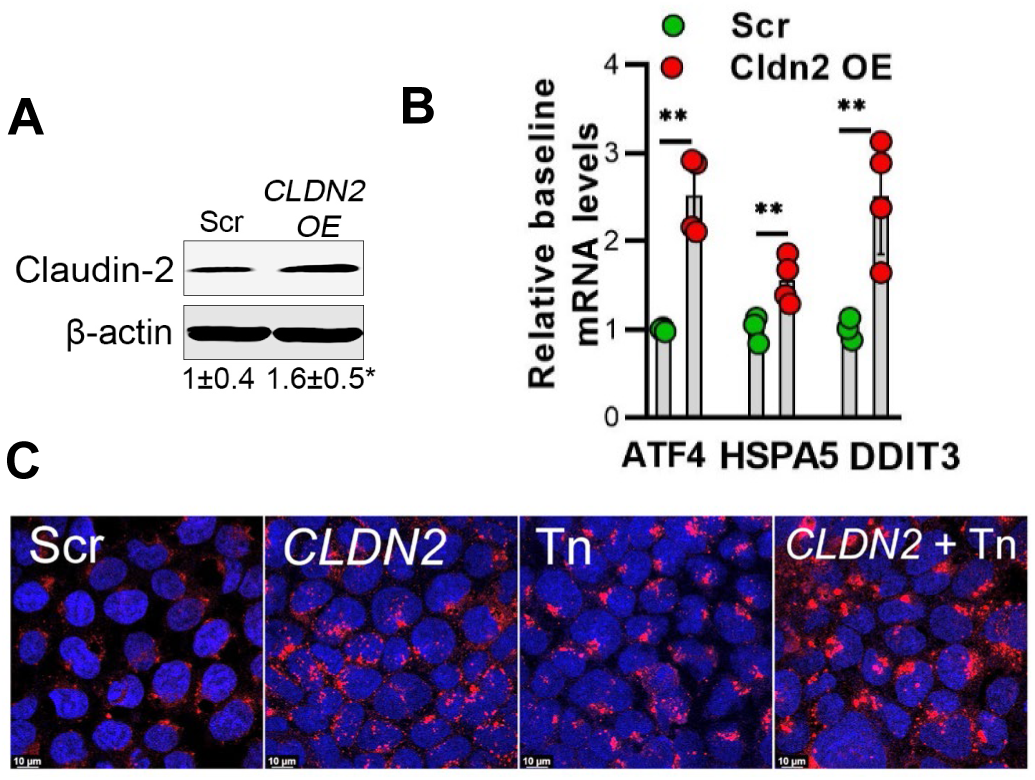
Claudin-2 over-expression (OE) induces ER stress and lysosomal damage. (A) Claudin-2 ORF in pCMV6-AC-GFP plasmid (and control plasmid) were used to transfect Caco-2 cells. *p< 0.03. (B) Cldn2 OE alone induced UPR genes ATF4, HSPA5 (Bip), and DDIT3 (CHOP) expression. **, p< 0.01. (C) Compared to the minimal cathepsin B staining (red) in scramble control cells, CLDN2 OE cells showed increased and diffuse staining for cathepsin B, comparable to tunicamycin-treated cells. White bar = 10μm.

### IRE1α kinase-mediated activation of AAK and inhibition of autophagy

To further evaluate the role of IRE1α kinase in AAK1 activation and regulation of autophagy, we prepared *IRE1α ^K599AΔ^* mutant in which the conserved lysine residue at 599 in the ATP binding site was substituted with alanine. As shown in **Fig. 8**, this mutation alone impaired the kinase activity of IRE1α (autophosphorylation of IRE1α, **Fig. 8A and B**), reduced phosphorylation of AAK1 target pAP2M1 levels (**Fig. 8A and C**). Also, importantly, baseline autophagy was increased after *IRE1α ^K599AΔ^* mutation (reduction in p62 levels, **Fig. 8A and D**), indicating significant role of IRE1α kinase in inhibition of autophagy. We also used NanoBRET intracellular kinase assay based on NanoLuc®-AAK1 kinase fusion vector to assess the role of IRE1α kinase in AAK1 activity. Compared to control scr cells, *IRE1Δ* cells showed multifold increase in AAK1 kinase BRET ratio (reduced intracellular AAK1 kinase engagement), indicating that in the absence of IRE1α, AAK1 activation is attenuated. This suggests that IRE1α interacts with and activates AAK1 (**Fig. 8E**).

**Fig. 8.**
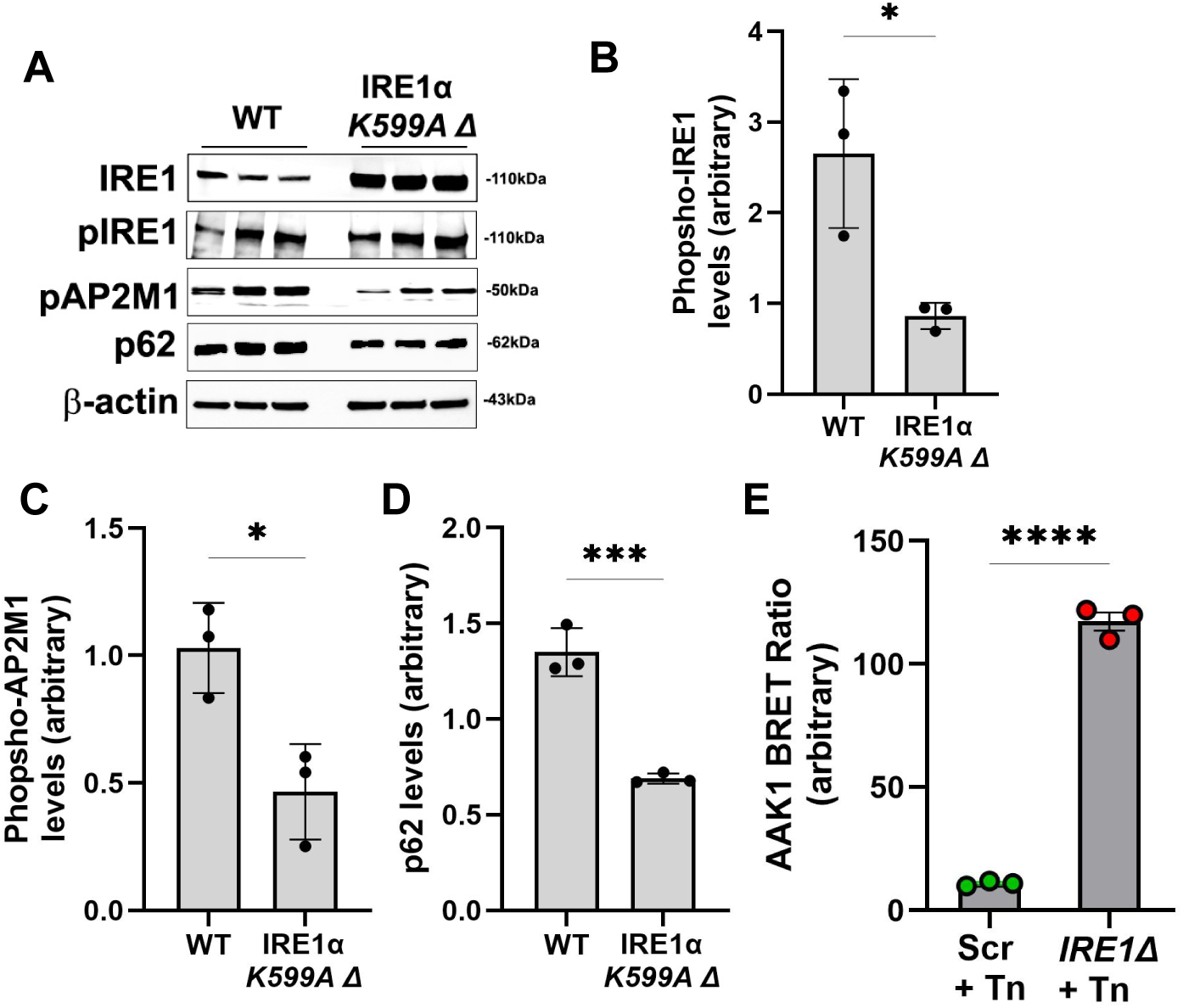
AAK1 regulation by IRE1 during the ER stress. (A) IRE1α kinase mutation (K599A) reduced baseline autophosphorylation of IRE1, phospho-AP2M1 levels, as well as p62 levels by ∼50% in Caco-2 cells. (B) Densitometry for phospho-IRE1 levels, normalized to total IRE1 from panel A. *p<0.05. (C) Densitometry for phosphor-AP2M1 levels, normalized to β-actin, from panel A. *p<0.05. (D) Densitometry for p62 levels, normalized to β-actin, from panel A. ***p<0.01. (E) In NanoBRET intracellular kinase assay, tunicamycin (Tn) treatment increases AAK1 BRET ratio by multi-fold in IRE1 KO (IRE1Δ) cells, indicating reduced intracellular AAK1 kinase engagement, which suggests that IRE1 directly interacts with AAK1 kinase domain and promotes its activation. ****, p<0.0001.

### Autophagy inhibits ER stress in-vivo

Finally, we investigated the protective role of autophagy against ER stress in murine colon using an autophagy-inducing, enterically delivered rapamycin (eRapa) diet. Tunicamycin administration increased protein levels of BiP and CHOP, while eRapa diet reduced tunicamycin-induced increase in ER proteins compared to vehicle diet (**Fig. 9A, B, and C**). This was also true under inflammatory condition. The mouse colonic levels of BiP and CHOP were markedly increased in chronic DSS colitis/ vehicle diet group. Administration of eRapa diet, however, reduced colonic BiP protein levels in chronic DSS colitis (**Fig. 9D and E**). Also, colonic p62 levels were increased in chronic DSS colitis, indicating disruption of autophagy during chronic inflammation. eRapa diet feeding during chronic DSS colitis promoted autophagy, as indicated by reduction in p62 levels (**Fig. 9D and F**). This data clearly shows that autophagy inhibits ER stress, at baseline as well as during chronic inflammation.

**Fig. 9.**
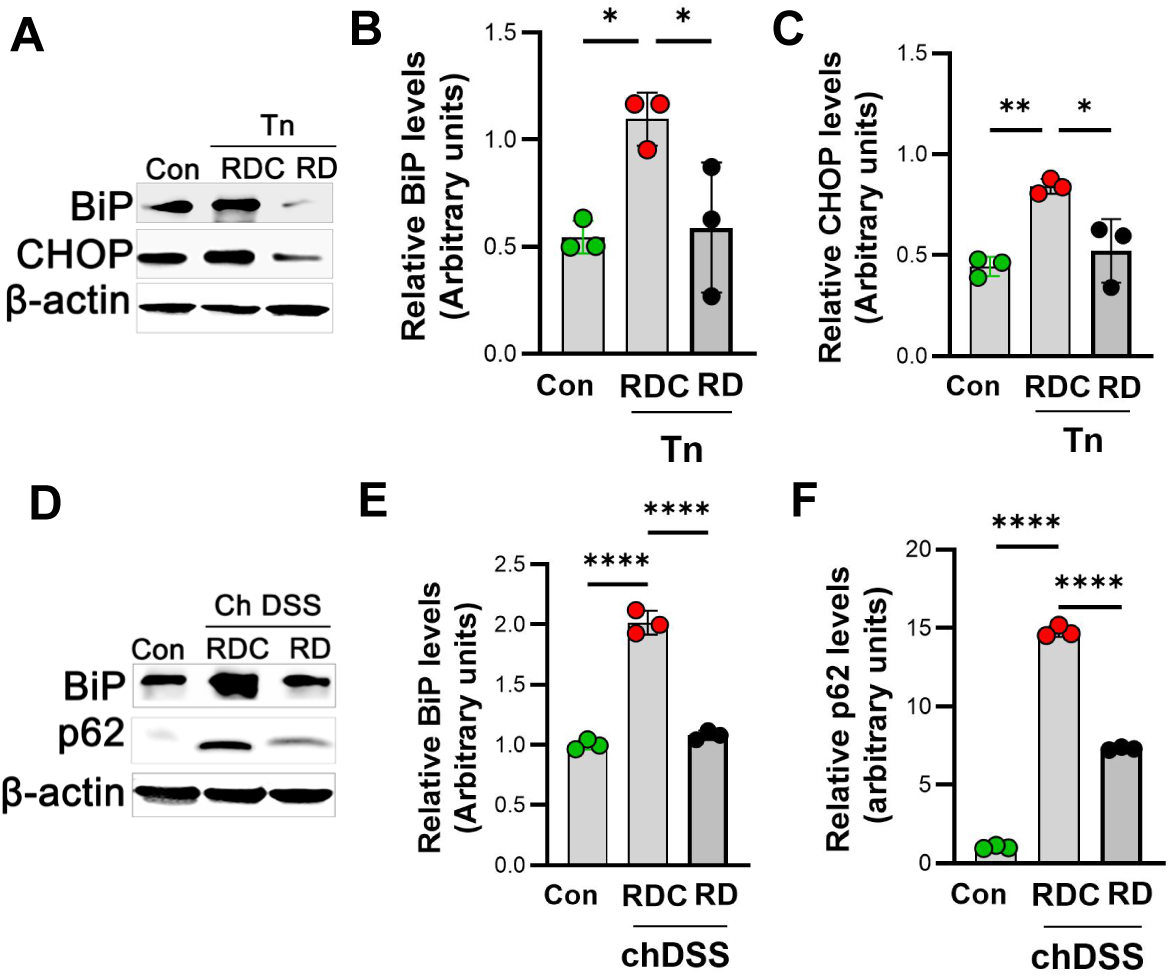
Rapamycin diet reduces ER stress in mouse colon. (A) Autophagy-inducing rapamycin diet (RD) reduced tunicamycin (Tn)-induced increase in UPR markers BiP and CHOP levels in mouse colon. Tn, 1mg/kg, i.p., 48 hours. RDC: Vehicle control diet. (B) Densitometry for BiP levels from panel A. (C) Densitometry for CHOP levels from panel A. (D) RD administration also reduced BiP levels in chronic DSS colitis (ch DSS) model and promoted autophagy (reduced p62 levels) in ch DSS colitis model. RD was given during the 3^rd^ round of ch DSS colitis model. Representation of 3 blots. All protein levels were normalized to β-actin levels. *p<0.05, **, p< 0.01 ***, p< 0.001, ****, p< 0.0001, One-way ANOVA.

## Discussion

The ER stress is one of the increasingly recognized signaling pathways involved in regulating intestinal homeostasis. Diverse developmental, infectious, metabolic, and pathophysiologic processes lead to accumulation of misfolded proteins and induce ER stress^31, 32^. The ER stress is followed by the unfolded protein response (UPR) aimed at restoration of cellular homeostasis^55, 56^. Persistent, unmitigated ER stress, however, overwhelms UPR and disrupts intestinal cellular homeostasis^57, 58^. Genes involved in the UPR as well as autophagy are known to be strongly associated with the risk for IBD development^40, 59^. Though ER stress is known to induce TJ barrier disruption^60–62^, the underlying mechanisms are not completely clear.

The interplay between autophagy and ER stress is a pivotal mechanism for epithelial barrier breach in IBD^63^. Using in-vitro cell culture, mouse studies, and human tissue, we showed that ER stress causes reduction in TER and increases paracellular permeability, accompanied by increased levels of pore-forming claudin-2 and disrupting the localization of barrier forming occludin. Furthermore, disruption of intestinal epithelial autophagy increases ER stress and associated TJ barrier dysfunction while induction of autophagy rescues ER stress-induced TJ barrier loss and cell survival. Thus, autophagy mitigates ER stress and ER stress-mediated disruption of TJ barrier. Our findings are consistent with previously shown role of autophagy in prevention of ER stress-induced blood-spinal cord barrier ^64, 65^.

The TJ barrier is regulated in several ways including gene expression, trafficking, and post-translational modifications^66^. We have previously showed that the AAK1-mediated activation of adaptor protein (AP2) μ subunit AP2M1, plays a crucial role in anchoring membrane claudin-2 to clathrin and LC3, for ultimate autophagic degradation.^10^ In this study, we demonstrated IRE1α as a relevant ER signaling pathway for the loss of TJ barrier during ER stress. Furthermore, IRE1α caused TJ disruption via AAK1. Significant reduction in AAK1 kinase engagement in IRE1α knock out cells during ER stress and prevention of phosphorylation of a well-known AAK1 target, AP2M1, by IRE1α kinase mutation supports IRE1α-mediated activation of AAK1. Thus, it appears that during initial ER stress, autophagy-mediated activation of AAK1 leads to endocytosis and cytoplasmic trafficking of claudin-2 for degradation, however, during persistent ER stress, loss of autophagic degradation leads to cytoplasmic aggregation of claudin-2. Also, it is remarkable that claudin-2 over-expression induces ER stress and causes lysosomal damage, further aggravating loss of claudin-2 homeostasis. Presently, the upstream pathway for AAK1 activation is unknown and the direct link between IRE1α-mediated activation of AAK1 is still unclear. However, AAK1 has been shown to be a substrate for pathways such as NDR1/2 ^67^ and Wnt ^68^, and it is possible that activated IRE1α during ER stress increases AAK1 activity through an intermediate mechanism that promotes membrane localization, phosphorylation, target proximity, or modulates inhibitory signaling pathways. On the other hand, sunitinib, an AAK-1 inhibitor, induced disruption of TJs, inhibited keratinocytes migration and proliferation, and triggered the intrinsic mitochondrial apoptotic pathway, which was reversed by inhibition of ER stress, suggesting a reciprocal regulation between AAK1 and ER stress.^69^

IRE1α, through its endoribonuclease activity of splicing XBP1 mRNA is essential for cellular homeostasis, and its total knockout increases susceptibility to ER stress ^40, 70^. However, IRE1α is also known to drive transmural ileitis phenocopying ileal CD in mice with Atg16l1 deletion in intestinal epithelial cells.^30^ Our study supports and provides an additional mechanism of TJ barrier disruption to the role of IRE1α in driving CD–like ileitis in the absence of active autophagy. Overall, using relevant cell culture and mouse models and validation in human explant cultures, we showed that activation of autophagy can ameliorate ER stress and associated intestinal TJ barrier disruption.

## Funding

This research work was supported in part by National Institute of Diabetes and Digestive and Kidney Diseases grant DK100562 (PN), DK114024 (PN). The authors also acknowledge support by the Peter and Marshia Carlino Fund for IBD Research.

## Acknowledgments

The authors thank the IBD and Colorectal Diseases Biobank, and Imaging (RRID:SCR_022526) and Animal Facility cores at the Penn State College of Medicine for their excellent technical assistance. The core services and instruments used in this project were funded, in part, by the Pennsylvania State University College of Medicine via the Office of the Vice Dean of Research and Graduate Students and the Pennsylvania Department of Health using Tobacco Settlement Funds (CURE). The content is solely the responsibility of the authors and does not necessarily represent the official views of the University or College of Medicine. The Pennsylvania Department of Health specifically disclaims responsibility for any analyses, interpretations or conclusions.

## Authors Contributions

PA and PN conceptualized and organized the study. PA, KS, AG and PN conducted experiments and analyzed the data. PA, KS, AG, AW and PN provided technical support.

PA, KS, AG, LH, GY and PN generated resources and provided scientific expertise. PA and PN drafted the manuscript.

## Disclosures

No conflicts of interest, financial or otherwise, are declared by the author(s).

## Data Availability Statement

The authors confirm that the data supporting the findings of this study are available within the article/ its supplementary materials.

## Notes

### Competing Interest Statement

The authors have declared no competing interest.

## References

[1] Mandel LJ, Bacallao R, Zampighi G: Uncoupling of the molecular ‘fence’ and paracellular ‘gate’ functions in epithelial tight junctions. Nature 1993, 361:552–5. 10.1038/361552a0

[2] Podolsky DK: Healing the epithelium: solving the problem from two sides. J Gastroenterol 1997, 32:122–6.

[3] Burden of digestive diseases in the United States, 2008. NIH Publication No. 09-6443. NIH, 2008.

[4] Ma TY, Anderson JM: Tight junction and intestinal barrier. Textbook of gastrointestinal physiology. Edited by Johnson LR. Elsevier Health Sciences, Philadelphia, PA, 2006. pp. 1559–94.

[5] Hollander D, Vadheim CM, Brettholz E, Petersen GM, Delahunty T, Rotter JI: Increased intestinal permeability in patients with Crohn’s disease and their relatives. A possible etiologic factor. Ann Intern Med 1986, 105:883–5.

[6] Turner JR: Intestinal mucosal barrier function in health and disease. Nature reviewsImmunology 2009, 9:799–809. 10.1038/nri2653

[7] Shen L, Weber CR, Raleigh DR, Yu D, Turner JR: Tight Junction Pore and Leak Pathways: A Dynamic Duo. Annu Rev Physiol 2010. 10.1146/annurev-physiol-012110-142150

[8] Van Itallie CM, Holmes J, Bridges A, Gookin JL, Coccaro MR, Proctor W, Colegio OR, Anderson JM: The density of small tight junction pores varies among cell types and is increased by expression of claudin-2. J Cell Sci 2008, 121:298–305. 10.1242/jcs.021485 [doi]

[9] Nighot P, Ma T: Role of autophagy in the regulation of epithelial cell junctions. Tissue Barriers 2016, 4:e1171284. 10.1080/21688370.2016.1171284

[10] Ganapathy AS, Saha K, Suchanec E, Singh V, Verma A, Yochum G, Koltun W, Nighot M, Ma T, Nighot P: AP2M1 mediates autophagy-induced CLDN2 (claudin 2) degradation through endocytosis and interaction with LC3 and reduces intestinal epithelial tight junction permeability. Autophagy 2021:1–18. 10.1080/15548627.2021.2016233

[11] Saha K, Ganapathy AS, Wang A, Morris NM, Suchanec E, Ding W, Yochum G, Koltun W, Nighot M, Ma T, Nighot P: Autophagy Reduces the Degradation and Promotes Membrane Localization of Occludin to Enhance the Intestinal Epithelial Tight Junction Barrier against Paracellular Macromolecule Flux. J Crohns Colitis 2022. 10.1093/ecco-jcc/jjac148

[12] Wong M, Ganapathy AS, Suchanec E, Laidler L, Ma T, Nighot P: Intestinal epithelial tight junction barrier regulation by autophagy-related protein ATG6/beclin 1. Am J Physiol Cell Physiol 2019, 316:C753–C65. PMC6580157. 10.1152/ajpcell.00246.2018

[13] Hampe J, Franke A, Rosenstiel P, Till A, Teuber M, Huse K, Albrecht M, Mayr G, De La Vega FM, Briggs J, Gunther S, Prescott NJ, Onnie CM, Hasler R, Sipos B, Folsch UR, Lengauer T, Platzer M, Mathew CG, Krawczak M, Schreiber S: A genome-wide association scan of nonsynonymous SNPs identifies a susceptibility variant for Crohn disease in ATG16L1. Nat Genet 2007, 39:207–11. 10.1038/ng1954

[14] Rioux JD, Xavier RJ, Taylor KD, Silverberg MS, Goyette P, Huett A, Green T, Kuballa P, Barmada MM, Datta LW, Shugart YY, Griffiths AM, Targan SR, Ippoliti AF, Bernard EJ, Mei L, Nicolae DL, Regueiro M, Schumm LP, Steinhart AH, Rotter JI, Duerr RH, Cho JH, Daly MJ, Brant SR: Genome-wide association study identifies new susceptibility loci for Crohn disease and implicates autophagy in disease pathogenesis. Nat Genet 2007, 39:596–604. 10.1038/ng2032

[15] Parkes M, Barrett JC, Prescott NJ, Tremelling M, Anderson CA, Fisher SA, Roberts RG, Nimmo ER, Cummings FR, Soars D, Drummond H, Lees CW, Khawaja SA, Bagnall R, Burke DA, Todhunter CE, Ahmad T, Onnie CM, McArdle W, Strachan D, Bethel G, Bryan C, Lewis CM, Deloukas P, Forbes A, Sanderson J, Jewell DP, Satsangi J, Mansfield JC, Wellcome Trust Case Control C, Cardon L, Mathew CG: Sequence variants in the autophagy gene IRGM and multiple other replicating loci contribute to Crohn’s disease susceptibility. Nat Genet 2007, 39:830–2. 10.1038/ng2061

[16] Saitoh T, Fujita N, Jang MH, Uematsu S, Yang BG, Satoh T, Omori H, Noda T, Yamamoto N, Komatsu M, Tanaka K, Kawai T, Tsujimura T, Takeuchi O, Yoshimori T, Akira S: Loss of the autophagy protein Atg16L1 enhances endotoxin-induced IL-1beta production. Nature 2008, 456:264–8. 10.1038/nature07383; 10.1038/nature07383

[17] Strisciuglio C, Duijvestein M, Verhaar AP, Vos AC, den Brink GR, Hommes DW, Wildenberg ME: Impaired autophagy leads to abnormal dendritic cell-epithelial cell interactions. J Crohns Colitis 2012. 10.1016/j.crohns.2012.08.009; 10.1016/j.crohns.2012.08.009

[18] Wildenberg ME, Vos AC, Wolfkamp SC, Duijvestein M, Verhaar AP, Te Velde AA, van den Brink GR, Hommes DW: Autophagy attenuates the adaptive immune response by destabilizing the immunologic synapse. Gastroenterology 2012, 142:1493–503.e6. 10.1053/j.gastro.2012.02.034; 10.1053/j.gastro.2012.02.034

[19] Cooney R, Baker J, Brain O, Danis B, Pichulik T, Allan P, Ferguson DJ, Campbell BJ, Jewell D, Simmons A: NOD2 stimulation induces autophagy in dendritic cells influencing bacterial handling and antigen presentation. Nat Med 2010, 16:90–7. 10.1038/nm.2069; 10.1038/nm.2069

[20] Wyatt J, Vogelsang H, Hubl W, Waldhoer T, Lochs H: Intestinal permeability and the prediction of relapse in Crohn’s disease. Lancet 1993, 341:1437–9.

[21] Arnott ID, Kingstone K, Ghosh S: Abnormal intestinal permeability predicts relapse in inactive Crohn disease. Scand J Gastroenterol 2000, 35:1163–9.

[22] Arrieta MC, Madsen K, Doyle J, Meddings J: Reducing small intestinal permeability attenuates colitis in the IL10 gene-deficient mouse. Gut 2009, 58:41–8. 10.1136/gut.2008.150888; 10.1136/gut.2008.150888

[23] Madsen KL, Doyle JS, Tavernini MM, Jewell LD, Rennie RP, Fedorak RN: Antibiotic therapy attenuates colitis in interleukin 10 gene-deficient mice. Gastroenterology 2000, 118:1094–105.

[24] Nighot PK, Hu CA, Ma TY: Autophagy enhances intestinal epithelial tight junction barrier function by targeting claudin-2 protein degradation. J Biol Chem 2015, 290:7234–46. PMC4358142. 10.1074/jbc.M114.597492

[25] Bertolotti A, Wang X, Novoa I, Jungreis R, Schlessinger K, Cho JH, West AB, Ron D: Increased sensitivity to dextran sodium sulfate colitis in IRE1beta-deficient mice. J Clin Invest 2001, 107:585–93. PMC199427. 10.1172/jci11476

[26] Hetz C: The unfolded protein response: controlling cell fate decisions under ER stress and beyond. Nature Reviews Molecular Cell Biology 2012, 13:89–102. 10.1038/nrm3270

[27] Tsuru A, Fujimoto N, Takahashi S, Saito M, Nakamura D, Iwano M, Iwawaki T, Kadokura H, Ron D, Kohno K: Negative feedback by IRE1β optimizes mucin production in goblet cells. Proc Natl Acad Sci U S A 2013, 110:2864–9. PMC3581977. 10.1073/pnas.1212484110

[28] Urano F, Wang X, Bertolotti A, Zhang Y, Chung P, Harding HP, Ron D: Coupling of stress in the ER to activation of JNK protein kinases by transmembrane protein kinase IRE1. Science 2000, 287:664–6. 10.1126/science.287.5453.664

[29] Kruse KB, Brodsky JL, McCracken AA: Characterization of an ERAD gene as VPS30/ATG6 reveals two alternative and functionally distinct protein quality control pathways: one for soluble Z variant of human alpha-1 proteinase inhibitor (A1PiZ) and another for aggregates of A1PiZ. Mol Biol Cell 2006, 17:203–12. PMC1345659. 10.1091/mbc.e04-09-0779

[30] Tschurtschenthaler M, Adolph TE, Ashcroft JW, Niederreiter L, Bharti R, Saveljeva S, Bhattacharyya J, Flak MB, Shih DQ, Fuhler GM, Parkes M, Kohno K, Iwawaki T, Janneke van der Woude C, Harding HP, Smith AM, Peppelenbosch MP, Targan SR, Ron D, Rosenstiel P, Blumberg RS, Kaser A: Defective ATG16L1-mediated removal of IRE1α drives Crohn’s disease–like ileitis. J Exp Med 2017, 214:401–22. 10.1084/jem.20160791

[31] Oakes SA, Papa FR: The role of endoplasmic reticulum stress in human pathology. Annu Rev Pathol 2015, 10:173–94. PMC5568783. 10.1146/annurev-pathol-012513-104649

[32] Schwärzler J, Mayr L, Vich Vila A, Grabherr F, Niederreiter L, Philipp M, Grander C, Meyer M, Jukic A, Tröger S, Enrich B, Przysiecki N, Tschurtschenthaler M, Sommer F, Kronberger I, Koch J, Hilbe R, Hess MW, Oberhuber G, Sprung S, Ran Q, Koch R, Effenberger M, Kaneider NC, Wieser V, Keller MA, Weersma RK, Aden K, Rosenstiel P, Blumberg RS, Kaser A, Tilg H, Adolph TE: PUFA-Induced Metabolic Enteritis as a Fuel for Crohn’s Disease. Gastroenterology 2022, 162:1690–704. 10.1053/j.gastro.2022.01.004

[33] Lu P, Struijs MC, Mei J, Witte-Bouma J, Korteland-van Male AM, de Bruijn AC, van Goudoever JB, Renes IB: Endoplasmic reticulum stress, unfolded protein response and altered T cell differentiation in necrotizing enterocolitis. PLoS One 2013, 8:e78491. PMC3806824. 10.1371/journal.pone.0078491

[34] Shkoda A, Ruiz PA, Daniel H, Kim SC, Rogler G, Sartor RB, Haller D: Interleukin-10 blocked endoplasmic reticulum stress in intestinal epithelial cells: impact on chronic inflammation. Gastroenterology 2007, 132:190–207. 10.1053/j.gastro.2006.10.030

[35] Tréton X, Pédruzzi E, Cazals-Hatem D, Grodet A, Panis Y, Groyer A, Moreau R, Bouhnik Y, Daniel F, Ogier-Denis E: Altered endoplasmic reticulum stress affects translation in inactive colon tissue from patients with ulcerative colitis. Gastroenterology 2011, 141:1024–35. 10.1053/j.gastro.2011.05.033

[36] Heazlewood CK, Cook MC, Eri R, Price GR, Tauro SB, Taupin D, Thornton DJ, Png CW, Crockford TL, Cornall RJ, Adams R, Kato M, Nelms KA, Hong NA, Florin THJ, Goodnow CC, McGuckin MA: Aberrant Mucin Assembly in Mice Causes Endoplasmic Reticulum Stress and Spontaneous Inflammation Resembling Ulcerative Colitis. PLoS Med 2008, 5:e54. 10.1371/journal.pmed.0050054

[37] Namba T, Tanaka K, Ito Y, Ishihara T, Hoshino T, Gotoh T, Endo M, Sato K, Mizushima T: Positive role of CCAAT/enhancer-binding protein homologous protein, a transcription factor involved in the endoplasmic reticulum stress response in the development of colitis. Am J Pathol 2009, 174:1786–98. PMC2671267. 10.2353/ajpath.2009.080864

[38] Hino K, Saito A, Asada R, Kanemoto S, Imaizumi K: Increased susceptibility to dextran sulfate sodium-induced colitis in the endoplasmic reticulum stress transducer OASIS deficient mice. PLoS One 2014, 9:e88048. PMC3912207. 10.1371/journal.pone.0088048

[39] Cao SS, Zimmermann EM, Chuang BM, Song B, Nwokoye A, Wilkinson JE, Eaton KA, Kaufman RJ: The unfolded protein response and chemical chaperones reduce protein misfolding and colitis in mice. Gastroenterology 2013, 144:989–1000.e6. PMC3751190. 10.1053/j.gastro.2013.01.023

[40] Kaser A, Lee AH, Franke A, Glickman JN, Zeissig S, Tilg H, Nieuwenhuis EE, Higgins DE, Schreiber S, Glimcher LH, Blumberg RS: XBP1 links ER stress to intestinal inflammation and confers genetic risk for human inflammatory bowel disease. Cell 2008, 134:743–56. PMC2586148. 10.1016/j.cell.2008.07.021

[41] Laukens D, Devisscher L, Van den Bossche L, Hindryckx P, Vandenbroucke RE, Vandewynckel YP, Cuvelier C, Brinkman BM, Libert C, Vandenabeele P, De Vos M: Tauroursodeoxycholic acid inhibits experimental colitis by preventing early intestinal epithelial cell death. Lab Invest 2014, 94:1419–30. 10.1038/labinvest.2014.117

[42] Akiyama T, Oishi K, Wullaert A: Bifidobacteria Prevent Tunicamycin-Induced Endoplasmic Reticulum Stress and Subsequent Barrier Disruption in Human Intestinal Epithelial Caco-2 Monolayers. PLoS One 2016, 11:e0162448. PMC5017626. 10.1371/journal.pone.0162448

[43] Nighot PK, Blikslager AT: Chloride channel ClC-2 modulates tight junction barrier function via intracellular trafficking of occludin. American journal of physiologyCell physiology 2012, 302:C178–87. 10.1152/ajpcell.00072.2011

[44] Schneider CA, Rasband WS, Eliceiri KW: NIH Image to ImageJ: 25 years of image analysis. Nat Methods 2012, 9:671–5. PMC5554542. 10.1038/nmeth.2089

[45] Karsli-Uzunbas G, Guo JY, Price S, Teng X, Laddha SV, Khor S, Kalaany NY, Jacks T, Chan CS, Rabinowitz JD, White E: Autophagy is required for glucose homeostasis and lung tumor maintenance. Cancer Discov 2014, 4:914–27. 10.1158/2159-8290.CD-14-0363 [doi]

[46] Nighot P, Al-Sadi R, Rawat M, Guo S, Watterson DM, Ma T: Matrix metalloproteinase 9-induced increase in intestinal epithelial tight junction permeability contributes to the severity of experimental DSS colitis. Am J Physiol Gastrointest Liver Physiol 2015, 309:G988–97. PMC4683300. 10.1152/ajpgi.00256.2015

[47] Van Itallie CM, Mitic LL, Anderson JM: Claudin-2 forms homodimers and is a component of a high molecular weight protein complex. The Journal of biological chemistry 2010. 10.1074/jbc.M110.195578

[48] Teckman JH, Perlmutter DH: Retention of mutant alpha(1)-antitrypsin Z in endoplasmic reticulum is associated with an autophagic response. Am J Physiol Gastrointest Liver Physiol 2000, 279:G961–74. 10.1152/ajpgi.2000.279.5.G961

[49] Ravikumar B, Vacher C, Berger Z, Davies JE, Luo S, Oroz LG, Scaravilli F, Easton DF, Duden R, O’Kane CJ, Rubinsztein DC: Inhibition of mTOR induces autophagy and reduces toxicity of polyglutamine expansions in fly and mouse models of Huntington disease. Nat Genet 2004, 36:585–95. 10.1038/ng1362

[50] Komatsu M, Waguri S, Ueno T, Iwata J, Murata S, Tanida I, Ezaki J, Mizushima N, Ohsumi Y, Uchiyama Y, Kominami E, Tanaka K, Chiba T: Impairment of starvation-induced and constitutive autophagy in Atg7-deficient mice. The Journal of cell biology 2005, 169:425–34. jcb.200412022 [pii]

[51] Walczak M, Martens S: Dissecting the role of the Atg12-Atg5-Atg16 complex during autophagosome formation. Autophagy 2013, 9:424–5. 10.4161/auto.22931 [doi]

[52] Ogata M, Hino S, Saito A, Morikawa K, Kondo S, Kanemoto S, Murakami T, Taniguchi M, Tanii I, Yoshinaga K, Shiosaka S, Hammarback JA, Urano F, Imaizumi K: Autophagy is activated for cell survival after endoplasmic reticulum stress. Mol Cell Biol 2006, 26:9220–31. PMC1698520. 10.1128/mcb.01453-06

[53] Shi B, Conner SD, Liu J: Dysfunction of endocytic kinase AAK1 in ALS. Int J Mol Sci 2014, 15:22918–32. PMC4284746. 10.3390/ijms151222918

[54] Lee H, Noh JY, Oh Y, Kim Y, Chang JW, Chung CW, Lee ST, Kim M, Ryu H, Jung YK: IRE1 plays an essential role in ER stress-mediated aggregation of mutant huntingtin via the inhibition of autophagy flux. Hum Mol Genet 2012, 21:101–14. 10.1093/hmg/ddr445

[55] Chakrabarti A, Chen AW, Varner JD: A review of the mammalian unfolded protein response. Biotechnol Bioeng 2011, 108:2777–93. PMC3193940. 10.1002/bit.23282

[56] Cláudio N, Dalet A, Gatti E, Pierre P: Mapping the crossroads of immune activation and cellular stress response pathways. EMBO J 2013, 32:1214–24. PMC3642686. 10.1038/emboj.2013.80

[57] Hiramatsu N, Chiang WC, Kurt TD, Sigurdson CJ, Lin JH: Multiple Mechanisms of Unfolded Protein Response-Induced Cell Death. Am J Pathol 2015, 185:1800–8. PMC4484218. 10.1016/j.ajpath.2015.03.009

[58] Hosomi S, Kaser A, Blumberg RS: Role of endoplasmic reticulum stress and autophagy as interlinking pathways in the pathogenesis of inflammatory bowel disease. Curr Opin Gastroenterol 2015, 31:81–8. PMC4592163. 10.1097/mog.0000000000000144

[59] Hooper KM, Barlow PG, Henderson P, Stevens C: Interactions Between Autophagy and the Unfolded Protein Response: Implications for Inflammatory Bowel Disease. Inflamm Bowel Dis 2019, 25:661–71. 10.1093/ibd/izy380

[60] Li B, Lee C, Chuslip S, Lee D, Biouss G, Wu R, Koike Y, Miyake H, Ip W, Gonska T, Pierro A: Intestinal epithelial tight junctions and permeability can be rescued through the regulation of endoplasmic reticulum stress by amniotic fluid stem cells during necrotizing enterocolitis. FASEB J 2021, 35:e21265. 10.1096/fj.202001426R

[61] Huang D, Xiong M, Xu X, Wu X, Xu J, Cai X, Lu L, Zhou H: Bile acids elevated by high-fat feeding induce endoplasmic reticulum stress in intestinal stem cells and contribute to mucosal barrier damage. Biochem Biophys Res Commun 2020, 529:289–95. 10.1016/j.bbrc.2020.05.226

[62] Cui YJ, Chen LY, Zhou X, Tang ZN, Wang C, Wang HF: Heat stress induced IPEC-J2 cells barrier dysfunction through endoplasmic reticulum stress mediated apoptosis by p-eif2α/CHOP pathway. J Cell Physiol 2022, 237:1389–405. 10.1002/jcp.30603

[63] Foerster EG, Mukherjee T, Cabral-Fernandes L, Rocha JDB, Girardin SE, Philpott DJ: How autophagy controls the intestinal epithelial barrier. Autophagy 2022, 18:86–103. PMC8865220. 10.1080/15548627.2021.1909406

[64] Zhou Y, Zhang H, Zheng B, Ye L, Zhu S, Johnson NR, Wang Z, Wei X, Chen D, Cao G, Fu X, Li X, Xu HZ, Xiao J: Retinoic Acid Induced-Autophagic Flux Inhibits ER-Stress Dependent Apoptosis and Prevents Disruption of Blood-Spinal Cord Barrier after Spinal Cord Injury. Int J Biol Sci 2016, 12:87–99. PMC4679401. 10.7150/ijbs.13229

[65] Zhou Y, Wu Y, Liu Y, He Z, Zou S, Wang Q, Li J, Zheng Z, Chen J, Wu F, Gong F, Zhang H, Xu H, Xiao J: The cross-talk between autophagy and endoplasmic reticulum stress in blood-spinal cord barrier disruption after spinal cord injury. Oncotarget 2017, 8:1688–702. PMC5352089. 10.18632/oncotarget.13777

[66] Nighot P, Ma T: Endocytosis of Intestinal Tight Junction Proteins: In Time and Space. Inflamm Bowel Dis 2021, 27:283-90. PMC7813749. 10.1093/ibd/izaa141

[67] Ultanir SK, Hertz NT, Li G, Ge WP, Burlingame AL, Pleasure SJ, Shokat KM, Jan LY, Jan YN: Chemical genetic identification of NDR1/2 kinase substrates AAK1 and Rabin8 Uncovers their roles in dendrite arborization and spine development. Neuron 2012, 73:1127–42. PMC3333840. 10.1016/j.neuron.2012.01.019

[68] Agajanian MJ, Walker MP, Axtman AD, Ruela-de-Sousa RR, Serafin DS, Rabinowitz AD, Graham DM, Ryan MB, Tamir T, Nakamichi Y, Gammons MV, Bennett JM, Couñago RM, Drewry DH, Elkins JM, Gileadi C, Gileadi O, Godoi PH, Kapadia N, Müller S, Santiago AS, Sorrell FJ, Wells CI, Fedorov O, Willson TM, Zuercher WJ, Major MB: WNT Activates the AAK1 Kinase to Promote Clathrin-Mediated Endocytosis of LRP6 and Establish a Negative Feedback Loop. Cell Rep 2019, 26:79–93.e8. PMC6315376. 10.1016/j.celrep.2018.12.023

[69] Wang J, Shen L, Chen S, Wang X, He Y, Zhang Y: Sunitinib Impairs Oral Mucosal Healing Through Endoplasmic Reticulum Stress-Mediated Keratinocyte Dysfunction. Cells 2025, 15. PMC12784725. 10.3390/cells15010001

[70] Zhang HS, Chen Y, Fan L, Xi QL, Wu GH, Li XX, Yuan TL, He SQ, Yu Y, Shao ML, Liu Y, Bai CG, Ling ZQ, Li M, Liu Y, Fang J: The Endoplasmic Reticulum Stress Sensor IRE1α in Intestinal Epithelial Cells Is Essential for Protecting against Colitis. J Biol Chem 2015, 290:15327–36. PMC4463471. 10.1074/jbc.M114.633560

